## Supplementary material for "Noncoding RNAs evolutionarily extend animal lifespan": methods and materials

### Computational environment

All computations were performed under Linux with Python 3.9 and R 4.3. Data analysis, including Loess and linear regression, significance tests, were done by existing R packages.

### Animal species data

The lifespan data of the animal kingdom was downloaded from AnAge Database of Animal Ageing and Longevity[1]. This includes 4215 animal species after filtering out 3 fungi and 1 plantae sample.

Genome info of these 4215 species were downloaded from NCBI database, including the latest sequence in fasta and annotation in GTF file format. Datasets, a command line tool developed by NCBI (https://www.ncbi.nlm.nih.gov/datasets/docs/v2/reference-docs/command-line/datasets/), was used to obtain data via the following command,

datasets download genome taxon ID --annotated --assembly-version latest --assembly-source RefSeq --include gtf,genome.

After downloading, only 333 out of 4215 samples had the complete genome and annotation files. These 333 data samples (Table_S1) were used for downstream analyses.

### Noncoding and protein length

Noncoding RNAs were derived from the GTF annotation files as a subset of the GTF annotation file by filtering out protein-coding genes.

Genome length represents the length of all chromosomes. Noncoding RNA and protein lengths for each sample were calculated by adding up all annotated noncoding RNA and protein lengths unless in figure 4A. Figure 4A counts motifs in all noncoding regions, including annotations and unannotated regions. The relative length of annotated noncoding RNAs and proteins was their lengths normalized by the corresponding genome length.

Among animal families, the length mean of annotated noncoding RNA and protein was calculated.

Gene number was the sum of all gene numbers in the corresponding category

**Motif frequency**

A total of 22868 motifs were generated from the complete permutation of 4 bases (ATCG) from 1 to 7 bases ($4^1+4^2+4^3+4^4+4^5+4^6+4^7$).

Motif frequency refers to the number of times a ncRNA motif appears in a sample. It was counted by mapping the motif to the sequence of an annotated ncRNA gene or a given ncRNA region. Then, add the count of all genes together to a total representing the frequency of this motif for these given samples.

**Zscore**

Zscore=(dat-mean(dat))/sd(dat)

**Lifespan motif inference**

A motif frequency matrix was constructed with 333 rows denoting 333 samples and 22868 columns as motifs. This matrix was used to find endogenous longevity motifs regulating lifespan using our software, FINET[2], as previously reported[3,4]. Briefly, FINET treats lifespan as a target (Y) in the matrix and searches for its regulators from the rest of motifs (X) via the elastic-net model. Elastic-net could produce more than 90% positive results[2]. To minimize errors, FINET introduces stability-selection[5] and randomly splits samples into m sub-groups (m = 2 in this study) and then searches for target-regulator interactions from each sub-group. If an interaction consistently occurs in m sub-groups, the type I error is very low and this error dramatically reduces when m values become large in complex biological data[2]. This stability selection repeats n times (n = 100 in this study). A frequency score (frequency in m*n trials) was calculated for each target-regulator interaction. A perfect frequency score (frequency score = 1) represents that an interaction always occurs (100%) in m*n random trials. This study used the frequency score of 0.60 (p = 0.60 as shown in the running command line below) as a cutoff to filter out interactions. The leftover interactions were treated as endogenous motifs interacting over lifespan independent of conditions.

We ran FINET as: julia finet.jl -c 120 -k 5 -n 100 -m 2 -a 0.5 -p 0.60 -i mydata.txt -o mynetwork and the result was reported here.

**Human normal tissue RNAseq data**

We downloaded a data set including 27 human normal tissues and 171 samples (Table_S2) published by L. Fagerberg et al[6]. The data set was in fastq format from SRA (Sequence Read Archive) with #PRJEB4337.

**Alignment and TPM calculation**

All RNA-seq files from SRA in fastq format were aligned to GRCh38.p13 and the read depth for 62,688 unique genes were counted by using STAR-2.5[7], with the following settings: runThreadN 30 --genomeDir GRCh38.p13.v43 --outSAMtype BAM Unsorted SortedByCoordinate --outFilterMultimapNmax 20 --outFilterType BySJout --chimSegmentMin 20 --alignSJoverhangMin 8 --alignSJDBoverhangMin 1 --quantMode TranscriptomeSAM GeneCounts --outFilterIntronMotifs RemoveNoncanonical --twopassMode Basic.

TPM was calculated as follows:

TPM = ratio / sum(ratio) * 1,000,000

Ratio = read counts / gene lengths

Log2(TPM) was used as the expression unit in the entire study.

The gene length was defined by the GENCODE[8] project in the GRCh38.p13.v43 GTF file.

**Mouse RNAseq and tissue**

A total of 1220 mouse RNA_seq samples were downloaded from ENCODE project[9], including 16 tissues and all types of RNA_seq samples. Those are all ENCODE's measurements so far. The RNA_seq processing was performed as the same way as done in human except setting the genome to mouse genome GRCm39.M31.

Reference:


1. Tacutu, R. *et al.* Human Ageing Genomic Resources: new and updated databases. *Nucleic Acids Res* **46**, D1083–D1090 (2018).
2. Wang, A. & Hai, R. FINET: Fast Inferring NETwork. *BMC Research Notes* **13**, 521 (2020).
3. Wang, A. Distinctive functional regime of endogenous lncRNAs in dark regions of human genome. *Computational and Structural Biotechnology Journal* **20**, 2381–2390 (2022).
4. Wang, A. Noncoding RNAs endogenously rule the cancerous regulatory realm while proteins govern the normal. *Computational and Structural Biotechnology Journal* **20**, 1935–1945 (2022).
5. Meinshausen, N. & Bühlmann, P. Stability selection. *Journal of the Royal Statistical Society: Series B (Statistical Methodology)* **72**, 417–473 (2010).
6. Fagerberg, L. *et al.* Analysis of the human tissue-specific expression by genome-wide integration of transcriptomics and antibody-based proteomics. *Mol Cell Proteomics* **13**, 397–406 (2014).
7. Dobin, A. *et al.* STAR: ultrafast universal RNA-seq aligner. *Bioinformatics* **29**, 15–21 (2013).
8. Harrow, J. *et al.* GENCODE: The reference human genome annotation for The ENCODE Project. *Genome Res* **22**, 1760–1774 (2012).
9. ENCODE Project Consortium. An integrated encyclopedia of DNA elements in the human genome. *Nature* **489**, 57–74 (2012).
